# NanoString nCounter® analysis of tissue microarray punch samples from colorectal carcinomas with invasive margins of the expansive and infiltrative type

**DOI:** 10.1101/2025.03.02.641055

**Authors:** Maja Hühns, Caterina Redwanz, Friedrich Prall

## Abstract

By a classical approach, the invasive margins of colorectal carcinomas can be typed as expansive or infiltrative, the latter portending a poor prognosis. Colorectal carcinomas with infiltrative-type invasive margins have long been known for their peculiar stroma and the relative absence of a lymphohistiocytic host response, but they have not been studied with contemporary gene expression techniques. Therefore, we submitted 18 microsatellite-stable colorectal carcinomas that could be typed as having invasive margins of the infiltrative (N = 7) or expansive type (N = 11) with confidence to nanoString nCounter® analysis (Tumor Signaling 360™ Panel). Tissue microarray punch samples were obtained from the tumors and prior to RNA extraction histological sections were prepared. The invasive margin types were mirrored surprisingly well in the two main clusters delineated by an unsupervised cluster analysis of the gene expression data, but tumor budding, the second type of colorectal carcinoma invasion phenotype, was not. Nanostring Annotation Scores were significant for signaling pathways (TGFb, PDGF, MET, FGFR), extracellular matrix remodeling, and anti-tumor immunity processes. However, any hopes that the tumour biology behind the two phenotypes of invasion could be pinpointed to differential expressions of a small set of genes were not fulfilled. Taken together, the data indeed give a molecular underpinning to the two invasion phenotypes, pointing out that matrix features and anti-tumor immunity are key. Nevertheless, we failed to gain a more detailed insight into the mechanics at work, and this may well be due to general limitations of the technology employed.

## Introduction

It is well appreciated in surgical pathology that, by histomorphology, the invasive margins of colorectal cancer differ between cases, and that these morphological features are quite effective in prognostication. Typing colorectal carcinomas’ invasive margins as expansive or infiltrative is the classical approach which was introduced by Jass et al. almost four decades ago [1]. Alleged issues with reproducibility [2] (which, however, can be alleviated if a dedicated effort is made [3]) may be one reason that these phenotypes of the invasive edge have largely fallen into oblivion. A second reason, probably more important, is widespread enthusiasm among surgical pathologists with tumor budding, the other morphological feature of invasive margins useful for prognostication [4], that by now is far outshining invasive margin typing.

No matter where a pathologist’s preference may lie, attention to phenotypes of the invasive margins or tumor budding, both, by looking down a microscope, let us catch a glimpse of tumor microenvironments, a hot topic in tumor biology. There have been a great number of studies designed to explore the tumor microenvironments at the invasive margins of colorectal carcinomas, and there is a long track list, dating back many years, of studies that applied immunohistochemistry in order to phenotype cells and matrix components in colorectal carcinomas putatively involved in tumor invasion. Tumor budding, in particular, has been studied intensively in this way (comprehensively reviewed in [5]). However, a study of the classical invasion phenotypes by the more recently available gene expression analysis techniques is outstanding.

The nanoString nCounter® technique is one of the methods available for large scale mRNA quantification, allowing simultaneous assessment of hundreds of genes in a single sample. The technique is robust because it dispenses with reverse transcription and DNA amplification, and it works well with RNA from archived paraffin blocks. Furthermore, there are available analysis tools that make the data amenable to researchers without a background in bioinformatics.

This study was conceived to explore if quantification of gene expression on a broad scale would provide any information on the tumor biology behind the phenomenon of the two types of invasive margins, *viz.* the infiltrative and the expansive type. Using the nanoString nCounter® technique on RNA extracted from tissue microarray punch samples that prior to the extraction had been submitted to histological study, we also attempted to correlate morphology with the molecular data.

## Materials and Methods

### Case selections and morphological studies

The colorectal carcinomas included in this study were accessioned to our institute per protocol of an ongoing colorectal carcinoma biobanking procedure which started in 2005 [6]. Patients’ written informed consent was obtained as part of the tumor banking procedures by collaborating doctors, and studies on the tumor material were approved by internal board review (Ethics Committee of Rostock University, ref. II HV 43/2004 and ref. A 2019-0187) which included approval that the senior author of the present study, by preparing the surgical pathology reports, had access to patient identifying information. Tissues used in this study were collected in the period between October 10^th^ 2019 and September 11^th^ 2023.

Surgical resections specimens were received fresh from the operating theatre and fixed overnight in buffered formalin (10%). None of the patients had received neoadjuvant radiation or chemotherapy. Dissections and reporting were done by the senior author. For this study, we selected 18 microsatellite-stable, standard-type adenocarcinomas of which the invasive margins could be typed as expansive or infiltrative with confidence. Tumor budding was assessed by a hot spot method on CK18 immunostains of whole tissue slides. Intratumoral tumor buds were counted on CK18 immunostains of the punch samples. Detailed information on the cases is given in Table 1.

**Table 1.** Clinicopathological data.

| Case ID | Invasive Margin-Type | Patient Data | Site | TNM | Tumor Budding | Buds in Punch (intratumoral budding) |
| --- | --- | --- | --- | --- | --- | --- |
| 70 | Expansive | m; 66 y | Rectum | G2 pT3 pN0 cM0 | Low | 0 |
| 75 | Expansive | f; 57 y | Sigmoid Colon | G2 pT3 pN0 cM0 | Low | 0 |
| 143 | Expansive | m; 83 y | Cecum | G2 pT4b pN0 cM0 | Low | 0 |
| 104 | Expansive | m; 84 y | Descending Colon | G2 pT2 pN0 cM0 | Low | 2 |
| 157 | Expansive | f; 64 y | Sigmoid Colon | G2 pT2 pN0 cM0 | Intermediate | 3 |
| 153 | Expansive | m; 51 y | Sigmoid Colon | G2 pT3 pN0 cM0 | Low | 4 |
| 79 | Expansive | m; 61 y | Transverse Colon | G2 pT3 pN1a cM1 | Low | 4 |
| 122 | Expansive | m; 59 y | Ascending Colon | G2 pT3 pN0 cM0 | High | 13 |
| 49 | Expansive | m; 54 y | Transverse Colon | G3 pT4b pN2b cM1 | Intermediate | 19 |
| 138 | Expansive | m; 54 y | Sigmoid Colon | G2 pT3 pN1b cM0 | High | 54 |
| 84 | Expansive | m; 73 y | Sigmoid Colon | G2 pT2 pN0 cM0 | High | 5 *) |
| 73 | Infiltrative | m; 61 y | Sigmoid Colon | G2 pT4a pN1c cM1 | High | 3 |
| 152 | Infiltrative | m; 81 y | Cecum | G2 pT3 pN0 cM0 | High | 6 |
| 149 | Infiltrative | m; 87 y | Sigmoid Colon | G2 pT4a pN1b cM0 | Intermediate | 8 |
| 141 | Infiltrative | f; 66 y | Descending Colon | G2 pT4a pN2b cM1 | Intermediate | 12 |
| 127 | Infiltrative | m; 55 y | Ascending Colon | G2 pT4a pN1a cM0 | High | 23 |
| 156 | Infiltrative | m; 63 y | Rectum | G2 pT4a pN2a cM1 | High | 33 |
| 145 | Infiltrative | f; 65 y | Sigmoid Colon | G2 pT4a pN1b cM1 | Intermediate | 38 |

\*) In this case, intratumoral budding was determined on the H&E section because CK18 immunohistochemistry was not available for technical reasons.

For each tumor, one paraffin-block that represented the tumor’s center was sampled with a 1.5 mm tissue microarrayer needle, and the punch samples were transferred into recipient paraffin-blocks for histological sections (H&E as well as CK18, CD8, and CD68 immunohistochemistry). Using QuPath [7], tumor areas and stroma areas in each punch sample were quantified on digital slides obtained from CK18 immunostains (pixel classifier), and digitized CD8 immunostains were used to quantify tumor cells and stromal cells as well as intratumoral T-cells and stromal T-cells (object classifier). In addition, stromal macrophage densities were assessed based on CD68 immunohistochemistry (pixel classifier). The remainders of the punch samples were then recovered from the paraffin-blocks for extraction of DNA and RNA.

### DNA and RNA isolation, microsatellite analyses, and panel NGS

RNA and DNA were extracted from the punch samples using the Maxwell® CSC RNA FFPE Kit and the Maxwell® CSC DNA FFPE Kit (Promega, Fitchburg, MA, USA), respectively, according to the manufacturer’s protocols. RNA and DNA concentrations were measured using the Quantus ™ Fluorometer (Promega).

Microsatellite analyses were done with the ESMO panel of microsatellite markers (Bat25, Bat26, NR-21, NR-24 and NR-27) and the quasi-monomorphic markers Cat25 and NR-22 [8, 9]. Furthermore, the DNA was used for targeted next-generation panel sequencing of genes frequently mutated in colorectal cancer. In this assay, a custom-designed NGS panel (AmpliSeq CRC v2 Panel; Illumina, San Diego, CA, USA) was used to sequence all exons and flanking intron regions of *ACVR1B*, *AMER1*, *APC*, *ARID1A*, *ATM*, *BRAF*, *CTNNB1*, *FBXW7*, *KRAS*, *NRAS*, *PIK3CA*, *SMAD2*, *SMAD4*, *SOX9*, *TCF7L2*, and *TP53*. Sequencing was performed on an Illumina iSeq100 with an input of 100 ng of genomic DNA. The variants were annotated using the Local Run Manager (Illumina) and the Integrative Genomics viewer (IGV version 2.8.9; Broad Institute) to determine the percentage of variant alleles (VAF).

### RNA-based expression analyses by nCounter technology

Gene expressions in the punch samples were studied with the nCounter® Tumor Signaling 360™ Panel (nanoString Technologies, WA, USA) which elegantly provides direct counts of mRNA molecules for each of the genes included in a panel. We here employed the Tumor Signaling 360™ Panel to profile the expression of 760 genes involved in tumor cell growth regulation, tumor microenvironment, and immune response.

A total of 100ng of RNA per sample was hybridized with capture probes and reporter probes using CodeSet hybridization at 65°C for 24 h. The hybridized samples were then processed on an automated nCounter® Prep Station, and the resulting nCounter® cartridges were analyzed using the nCounter® Digital Analyzer. Data were analyzed using ROSALIND® Analysis Software (Rosalind Inc., San Diego, CA, USA), with a HyperScale architecture developed by ROSALIND®. As part of the quality control step, read distribution percentages, violin plots, identity heat maps and sample MDS plots were generated. Normalization, fold changes and p-values were calculated using the criteria provided by nanoString. ROSALIND® follows the nCounter® Advanced Analysis protocol, where the counts within a lane are divided by the geometric mean of the normalization probes from the same lane. The housekeeping probes to be used for normalization are selected based on the geNorm algorithm implemented in the NormqPCR R library [10]. Fold changes and significance score (*p*-value) were calculated using the fast method as described in the nCounter^®^ Advanced Analysis 2.0 User Manual (nanoString). Significant *p*-values (*p* < 0.05) were adjusted for multiple genetic comparisons using the Benjamini– Hochberg method of estimating false discovery rates [11]. Clustering of genes for the final heatmap of differentially expressed genes was done using the Partitioning Around Medoids method and the fpc R library (version 2.2-13) [12]. All samples passed the quality control test. In comparing tumors with an invasive margin of the expansive-type versus the infiltrative-type, group changes in gene expression were presented as fold-changes. Differential expression of a gene was called at a threshold of greater than 1.5-fold change along with a *p*-value<0.05.

## Results

This study was made with 18 microsatellite-stable colorectal adenocarcinomas of the standard-type which were selected from our tumor bank to include tumors with a prototypical invasive margin of the expansive or infiltrative type (11 and 7 cases, respectively). We then built tissue microarrays (1.5 mm needle) from the tumors by sampling paraffin-blocks that represented the tumor centers. By taking H&E sections from the microarrays, this procedure provided good visual control over the tissues which went into RNA extractions and the nCounter assays. For illustration, microscopic images of one tumor with expansive-type invasive margins and one tumor with infiltrative-type invasive margins and the corresponding punch samples are shown in Figs 1 and 2, respectively. Furthermore, we obtained CK18, CD8, and CD68 immunohistochemistry to enhance the morphological assessments prior to RNA extraction from the punches after removal from the tissue microarrays.

**Fig 1.**
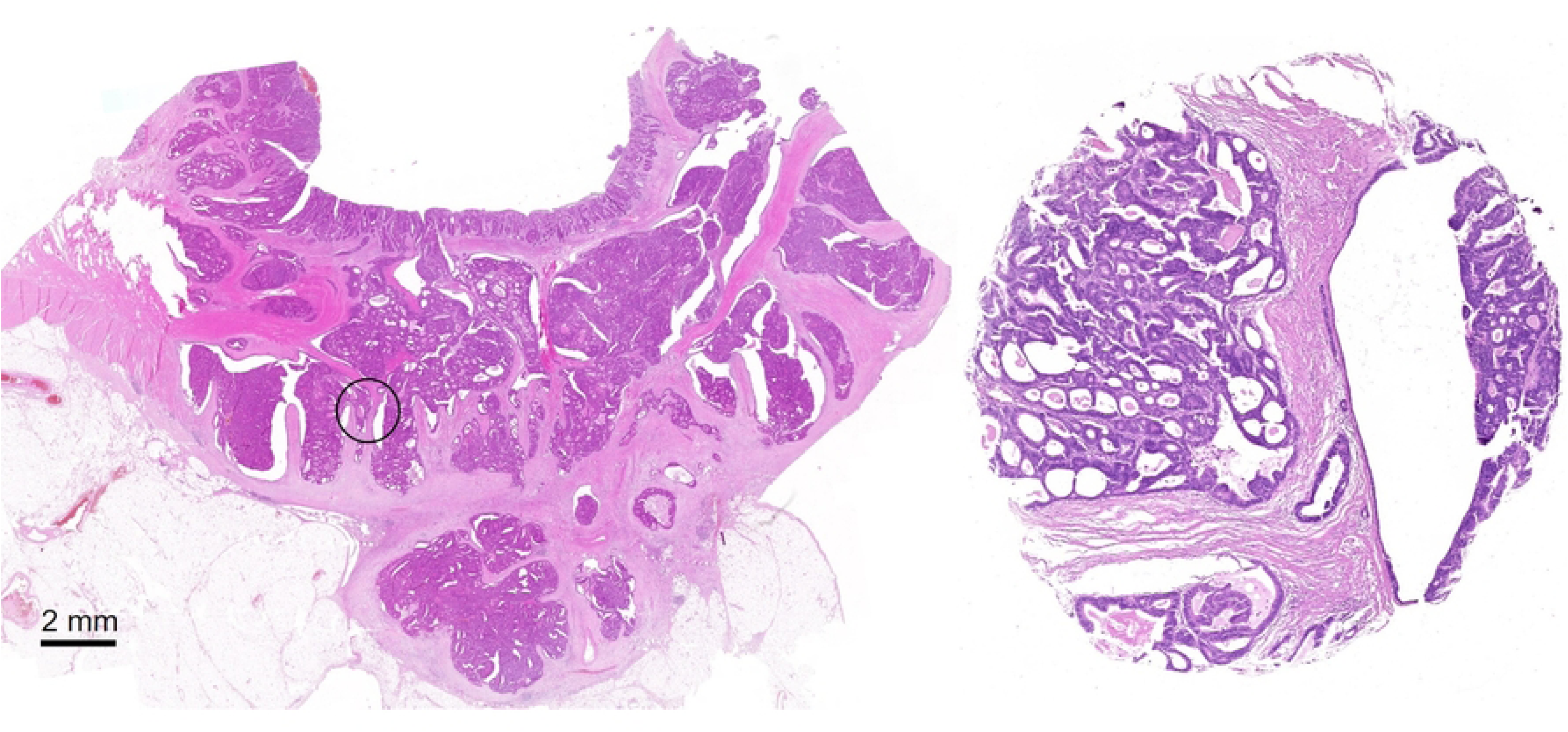
Colorectal cancer with an expansive-type invasive margin (case 75) and a microscopic image of the punch sample prior to RNA extraction. The site from which the punch sample was taken is marked on the whole mount slide (circle).

**Fig 2.**
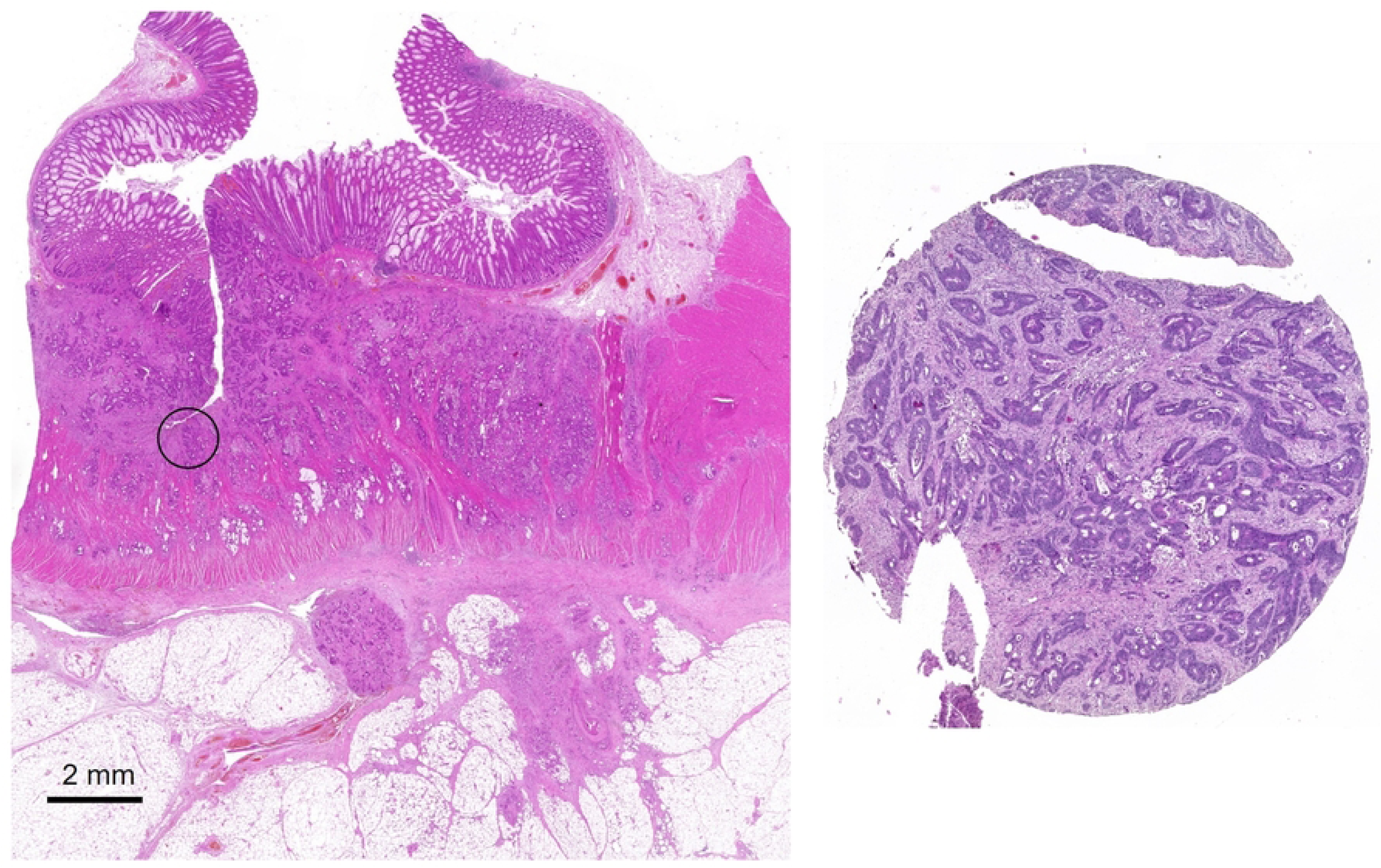
Colorectal cancer with an infiltrative-type invasive margin (case 156) and a microscopic image of the punch sample prior to RNA extraction. The site from which the punch sample was taken is marked on the whole mount slide (circle).

The nCounter assay worked well with the RNA extracted from the punch samples in the tissue microarrays. The gene expression data were submitted to unsupervised two-dimensional cluster analysis and heatmapping as implemented with ROSALIND™, which is shown in Fig 3. This analysis separated the 18 tumors into two main clusters: 9 tumors were assigned to Cluster 1, all of which had an expansive-type invasive margin; Cluster 2 comprised the remaining tumors, all the 7 cases with an infiltrative-type invasive margin and 2 expansive-types (cases 143 and 104). Thus, the classification of tumors by invasion phenotypes was fairly well mirrored in the gene expression data derived from the nCounter panel employed here; however, neither the degree of tumor budding nor intratumoral tumor budding was.

**Fig 3.**
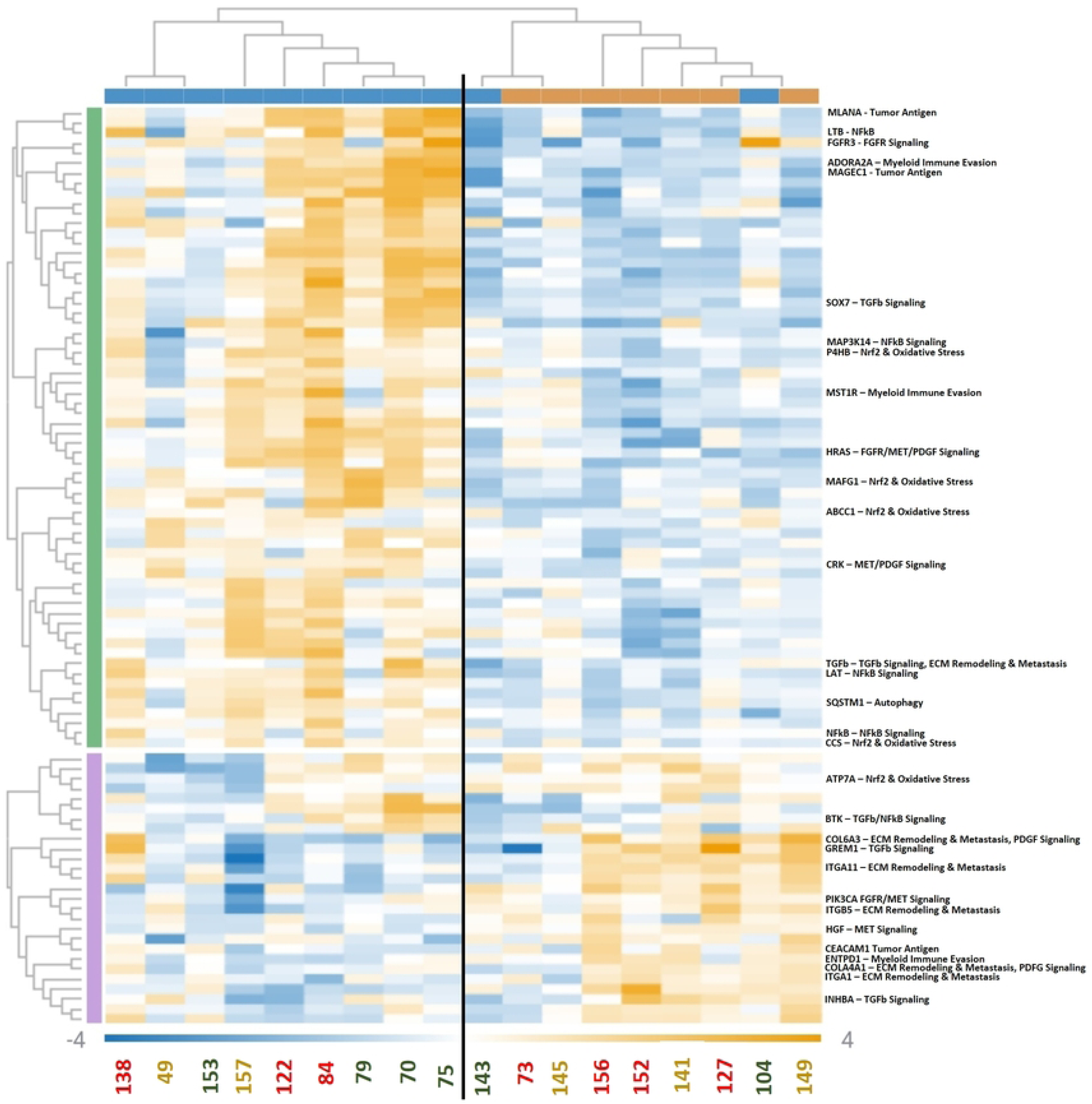
Heatmap of the nCounter® Tumor Signaling 360™ Panel data and two-dimensional cluster analysis (fold change of 1.5 up- or downregulated, p-value < 0.05). Cases are arranged in columns (blue bar at the top denotes expansive-type margin and yellow bar infiltrative-type margin) and genes are arranged horizontally (genes/pathways with significant scores are detailed to the right). Note two main clusters, one exclusively comprising tumors with expansive margins (cluster to the left) and another cluster which contains all the tumors with infiltrative-type invasive margins and two with expansive-type invasive margins (cluster to the right). IDs of the cases are as in Table 1, and the degree of tumor budding is colour-coded: red – high degree, yellow – intermediate degree, green – low (for intratumoral tumor budding see Table I).

Clustering of the cases was built on 91 differentially expressed genes of the 760 targets included in the nCounter® Tumor Signaling 360™ Panel. Pathway analysis as implemented with the ROSALIND™ suite (nanoString Annotations) identified differential expression of genes in two main categories of tumor cell biology. Specifically, nanoString Annotation Scores were significant for (i) genes involved in matrix receptor and growth factor signaling (TGFb signaling, PDGF signaling, MET signaling, FGFR signaling) as well as matrix turnover (extracellular matrix remodeling, ECM); and (ii) genes involved in tumor immunity (myeloid immune invasion, NFkB signaling, tumor antigen). Furthermore, nanoString Annotation Scores were significant in the category of autophagy and Nrf2 & Oxidative Stress. Genes and their respective nCounter® annotations are shown in Fig 3.

By a second approach to the data, we explored which genes were expressed differentially when comparing the two groups. Box-plots representing expression data for some of the genes with the greatest differences are shown in Fig 4 (see Fig. S1 for a complete set of data). As can be seen from these plots, while global differences were seen (and statistically significant when comparing the means), there always were cases which by gene expressions cut across the groups, and differential expression of any gene in all the cases was never observed.

**Fig 4.**
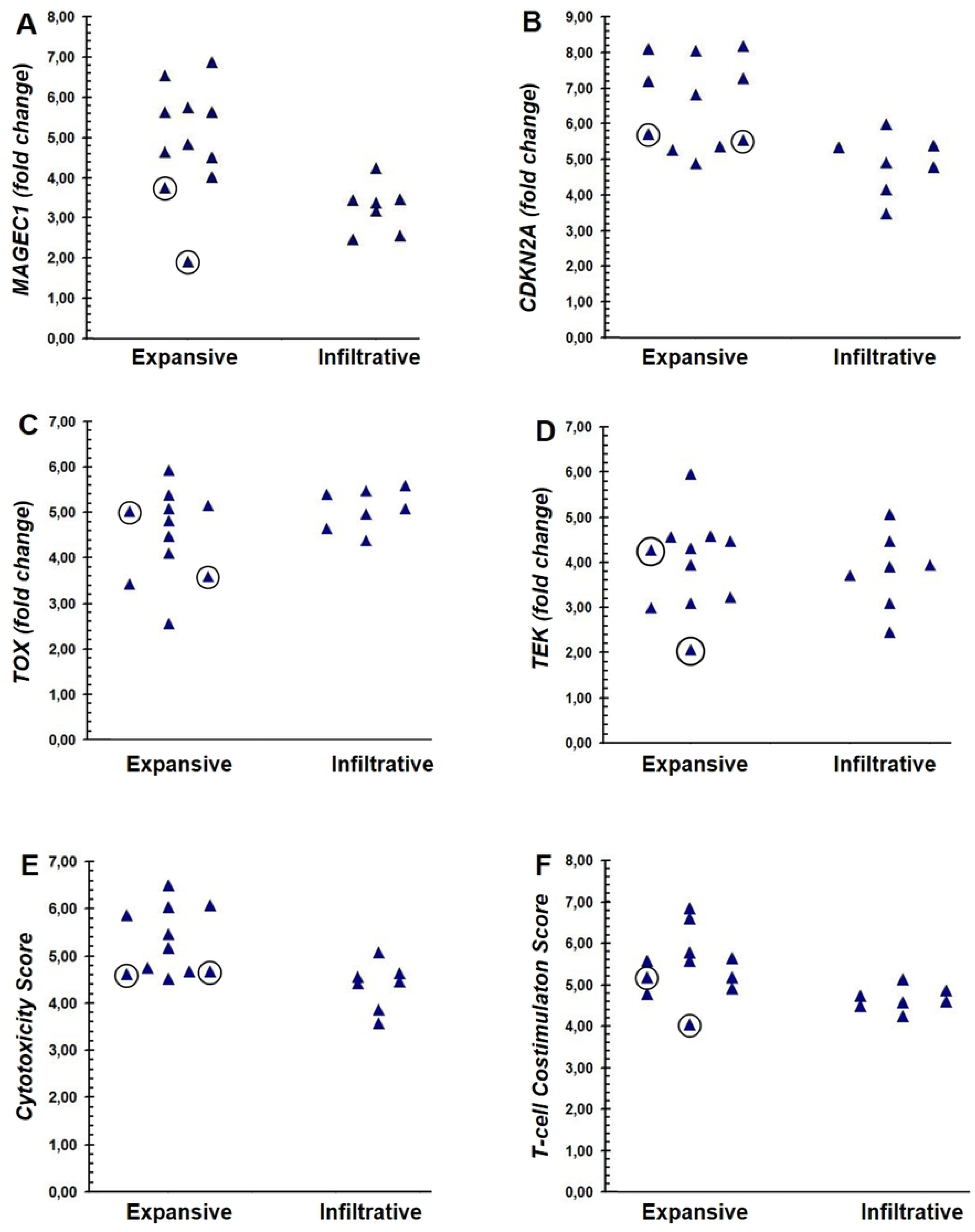
Box plots comparing nCounter data obtained for tumors with expansive-type and infiltrative-type invasive margins for four genes. (A – D, MAGEC1, CDKN2A, TOX, and TEC, respectively), and for the nCounter Cytotoxicity Score (E) and the T-cell Costimulation Score (F). Cases ID104 and ID143, which cut across the classes in the cluster analysis (cf. Fig 3), are encircled.

A quantitative evaluation of CD8 and CD68 immunohistochemistry showed that densities of T-cell or macrophage infiltration did not differ significantly when comparing the samples from tumors with invasive margins of expansive or infiltrative types. By contrast, the cytotoxicity score and the T-cell costimulation score, which are implemented with ROSALIND™ and combine expression data from several genes involved in these immunological processes, were clearly lower for the cases with infiltrative-type invasive margins (Fig 4).

Somatic mutations of the colorectal carcinomas included in this study were studied by panel NGS which included the genes most frequently mutated in microsatellite-stable tumors. Specifically, we addressed if any of the gene mutations interrogated by our panel would be more common in either the infiltrative-type or the expansive-type tumors. As can be seen in Table SI, this appeared not to be the case.

## Discussion

In this study, we addressed if gene expression profiles obtained with the nCounter Tumor Signaling 360™ Panel would give a clue to the biology behind the two different invasion phenotypes of colorectal carcinomas, namely the invasive margins of the expansive and infiltrative types. The panel we selected quantifies mRNA of 760 genes involved in a broad range of cellular processes taking place in tumor growth. Specifically, to name the most important with regard to colorectal cancer, the panel interrogates genes involved in signaling pathways that affect cell proliferation (MAPK, MET, PI3K, TGFb, Wnt, FGFR, EGFR, Hedgehog, ERBB2) and guarding of the genome (TP53, DNA damage repair); genes involved in cell adhesion and cell migration, epithelial-mesenchymal transition (EMT), and remodeling of the extracellular matrix; genes that promote inflammation (chemokine signaling and NF-kB signaling among others); and a number of genes that reflect immune processes (e.g. cytotoxicity, T-cell costimulation, and T-cell exhaustion). The mRNA submitted to the assays was extracted from punch samples that were taken from non-necrotic areas of the tumors away from the invasive margins proper, i.e. they did not contain the tumor-stroma interface at the deepest level of invasion. This is counterintuitive, at first sight, in a study that addresses invasive margins and aspects of its cellular biology. However, we had reasons for this approach: First, punch samples, regardless of their diameters (1.5 mm in our case) are circles, and this basically makes it impossible to obtain material that contains a well balanced proportion of invasive margin tumor cells and stroma. As was our experience in preliminary attempts, even if much care is taken, it usually is either an exceeding amount of tumor or an unacceptable preponderance of stroma. Second, the nanoString nCounter® technique does not allow distinction between mRNA derived from tumor cells and stromal cells, which makes it undesirable to submit samples with substantially different amounts of tumor and stromal cells. After some deliberations, these constraints let us decide to sample the tumors’ centers with a view to obtaining material which, at least roughly, offered similar tumor-stroma relations.

The gene expression data were used for hierarchical cluster analysis which delineated two main groups: a cluster exclusively made up of expansive-type margin tumors on one side (nine tumors), and a second cluster that contained all the seven infiltrative-type margin tumors and two of the expansive-types. Considering that the mRNA that went into the assay was extracted from punch samples which, as discussed above, were taken from the tumors’ centers and not the invasive margins *sensu strictu* this is somewhat remarkable. It may signify that cellular processes which lead to the different phenotypes of invasion are elaborated throughout the tumors and not restricted only to the invasive margins. While observable to pathologists by microscopic study only of whole tissue slides, mRNA profiles processed bioinformatically apparently can “see” the difference even in punch samples.

Notably, the degree of tumor budding, the second invasion phenotype of colorectal carcinomas did not impact on the cases’ assignments to clusters. This is not entirely unexpected because tumor budding essentially, at least by morphology, is an invasive margin feature, and therefore it is of prevalence in areas not specifically investigated by our approach. There also is intratumoral budding [13] which, however, also did not translate into the cases’ cluster assignments. The failure of tumor budding to impact appreciably the clustering results, whereas margin typing did, actually underscores that the distinction of expansive-type vs. infiltrative-type invasive margins reflects properties of colorectal carcinomas that are based on a cellular biology different from tumor budding. Notably in this respect, WNT signaling, a key player in tumor budding [14], was not found to differ in the two classes of invasive margins.

The clustering results obtained here (Fig 3) indeed strike the very themes of tumour biology which, from a morphological point of view, are expected to be played. Infiltrative-type colorectal cancers are noted for their peculiar matrix and for their immune-deserted phenotype [15], and this appears to be reflected in the pathways singled out by the nanoString Annotations, namely ECM remodeling, tumor antigen, and TGFb signaling (among other signalling pathways). However, any hopes that the tumour biology behind the two phenotypes of invasion could be pinpointed to differential expressions of a small set of genes were not fulfilled by our results. In fact, differences in gene expressions, though significant by the metrics, numerically were fairly slight. It is unlikely that they would be detectable in immunohistochemical studies.

Published studies that applied the nanoString nCounter® technology to colorectal carcinomas are limited to-date. The study by Chen et al. [16] compared the nanoString nCounter® technology with the Affymetrix U133 Plus 2.0 Gene Chip-Array to explore technical issues and also proposed a small set of genes as suitable for a prognostic nCounter assay. The study by Ragulan et al. [17] tested if the nanoString nCounter® technology would yield a classification of colorectal carcinomas similar to the Consensus classification [18] which is based on gene expression data generated by Affymetrix, Agilent or RNA sequencing assays. A large series of colon cancers stages II and III were submitted to nCounter analysis in a prognostic/predictive factor study by Xu et al. [19], and Yin et al. [20] applied the nanoStrings nCounter® PanCancer IO 360 panel to a series of colorectal cancers to explore the prognostic effect of a T-cell based inflammation signature on prognosis. Each study reported significant results. However, looking at the list of genes that are reported as being expressed differentially between the cases up for comparisons fosters a feeling of uneasiness: in each of these studies, the set of genes reported as significant is always a new one, begging the question why this would be so.

Attributing this to the assays’ imprecisions does not appear to be a convincing answer because the nanoString nCounter® assays provide very reproducible results upon repeat runs [17]. In side-by-side comparisons of the nanoString nCounter® and the Affymetrix technologies discrepancies are somewhat larger but nevertheless substantial. Much rather, we suspect, the extensive statistical processing of the data may be a reason, and the nature of the tissue samples that are entered into the gene expression assays most certainly are.

Gene expression studies generate very large datasets which could not be handled “by the naked eye” as academic pathologists did in the past with expression data obtained by e.g. one or a limited set of RT-PCR assays (or immunohistochemistry results). However, in the past, gene expression studies with tumor tissues were hypothesis-driven whereas contemporary large-scale gene expression studies basically are exploratory, sifting through hundreds or even thousands of data-points in search for interrelations. It is a well known drawback incurred by this approach that, no matter the p-values, results are difficult to reproduce with independent samples. While, without any doubt, the gene expression assays available today are fascinating (and would have felt to be science-fiction twenty years ago), a critical eye on the “black-box statistics” behind the shining results may be warranted.

Tissues are the second aspect. The tissues entered into large-scale gene expression studies usually are controlled for their tumor cell content. However, there never is any information on the tumor microenvironment of the study samples which, to name the most obvious aspects, is made up of the different cell types present in a sample and the extracellular matrix; the position within a tumor as a whole may also play a role, e.g. vicinity to (larger) areas of necrosis or pronounced granulocytic inflammation. Leaving these aspects up to chance (i.e. the random sampling of the tumours) and having no means to revisit morphology after the gene expression data have come in, may very well level out differences in gene expressions that are important in a microenvironment or impair reproducibility of results in different series.

The present study was made on a very modest set of cases because we included but 18 tumors, which is far less than in published studies. However, there are two aspects which may explain why, in spite of this limitation, the results were surprisingly clear-cut. First, this study was hypothesis-driven in that we tested if phenotypes of invasion would be mirrored in gene expressions. Second, we submitted RNA to the nanoString nCounter® assays from tumour tissues that prior to RNA extractions had been studied by histology, and this is an approach that, to the best of our knowledge, has not been tried before. We are well aware of the fact that the morphology-gene expression correlations presented here have not in the least been exhausted to its full extent by us. It may be worthwhile, for example, to apply artificial intelligence algorithms to the digital images of the punches, and immunohistochemistry could have been more extensive to characterize the different cells in the samples (i.e. the “players” in the different microenvironments). Indeed, it would be gratifying to us if our approach were to be an incentive to working groups with resources beyond ours. In conclusion, we have shown here that submitting tissue microarray punch samples to nanoString nCounter® analyses works well in morphologic-molecular correlations. This should be an approach that can be extended to the study of any kind of solid tumor. As regards the colorectal cancers studied here, the data indeed appear to give a molecular underpinning to the two invasion phenotypes. They point out that – not unexpectedly – matrix features and anti-tumor immunity are key. However, inspite of our honing in on the microenvironment as much as can be done short of single cell expression studies and yet more complex bioinformatics (the “next generation technology” coming up at the horizon), we failed to gain a more detailed insight into the mechanics at work, and this may well be due to limitations of the technology employed.

## Acknowledgements

Not applicable.

## Supporting information

**S1 Fig.** Gene expression data for all genes with fold-change expression ≥2 or ≤2.

**S1 Table.** Gene mutations found by NGS panel sequencing.

## References

[1] Jass JR, Atkin WS, Cuzcik J, Bussey HJ, Morson BC, Northover JM, Todd IP. The grading of rectal cancer: historical perspectives and a multivariate analysis of 447 cases. Histopathology. 1986;10:437 – 459.

[2] Jass JR, Allen JP, Chan YF. Assessment of invasive growth pattern and lymphocytic infiltration in colorectal cancer. Histopathology. 1996;28:543 – 548.

[3] Wöhlke M, Schiffmann L, Prall F. Aggressive colorectal carcinoma phenotypes of invasion can be assessed reproducibly and effectively predict poor survival: interobserver study and multivariate survival analysis of a prospectively collected series of 299 patients after potentially curative resections with long-term follow-up. Histopathology. 2011;59:857 – 866.

[4] Prall F. Tumour budding in colorectal carcinoma. Histopathology. 2007;50:151 – 162.

[5] Lugli A, Zlobec I, Berger MD, Kirsch R, Nagtegaal ID. Tumour budding in solid cancers. Nature Rev Clin Oncol. 2021;18:101 – 115.

[6] Mullins CS, Micheel B, Matschos S, Leuchter M, Bürtin F, Krohn M, et al. Integrated biobanking and tumor model establishment of human colorectal carcinoma provides excellent tools for preclinical research. Cancers. 2019;11:1520 - 1536.

[7] Bankhead P, Loughrey MB, Fernández JA, Dombrowski Y, McArt DG, Dunne PD, et al. QuPath: open source software for digital pathology image analysis. Sci Rep. 2017;7:e16878. 10.1038/s41598-017-17204-5.

[8] Luchini C, Bibeau F, Ligtenberg MJL, Singh N, Nottegar A, Bosse T, et al. ESMO recommendations on microsatellite instability testing for immunotherapy in cancer, and its relationship with PD-1/PD-L1 expression and tumour mutational burden: a systematic review-based approach. Ann Oncol. 2019;30:1232 – 1243.

[9] Ostwald C, Linnebacher M, Weirich V, Prall F. Chromosomally and microsatellite stable colorectal carcinomas without the CpG island methylator phenotype in a molecular classification. Int J Oncol. 2009;35:321 – 327.

[10] Perkins JR, Dawes JM, McMahon SB, Bennett DL, Orengo C, Kohl M. ReadqPCR and NormqPCR: R packages for the reading, quality checking and normalisation of RT-qPCR quantification cycle (Cq) data. BMC Genomics. 2012;13:296. 10.1186/1471-2164-13-296.

[11] Ferreira, J. A. (2007). The Benjamini-Hochberg method in the case of discrete test statistics. Int J Biostat. 2007;3 (1) Art.11. doi: 10.2202/1557-4679.1065.

[12] Hennig, C. (2019). Cran-package fpc. Vienna: R Core Team. https://cran.r-project.org/web/packages/fpc/index.html.

[13] Lugli A, Vlajnic T, Giger O, Karamitopoulou E, Patsouris ES, Peros G, et al. Intratumoral budding as a potential parameter of tumor progression in mismatch repair-proficient and mismatch repair-deficient colorectal cancer patients. Hum Pathol. 2011;42:1833 – 1840.

[14] Hlubek F, Brabletz T, Budczies J, Pfeiffer S, Jung A, Kirchner T. Heterogenous expression of WNT/b-catenin target genes within colorectal cancer. Int J Cancer. 2007;121:1941 – 1948.

[15] Ueno H, Shinto E, Shimazaki H, Kajiwara Y, Sueyama T, Yamamoto J, et al. Histologic categorization of desmoplastic reaction: its relevance to the colorectal cancer microenvironment and prognosis. Ann Surg Oncol. 2015;22:1504 – 1512.

[16] Chen X, Deane NG, Lewis KB, Li J, Zhu J, Washington MK, et al. Comparison of Nanostring nCounter Data on FFPE Colon Cancer Samples and Affymetrix Microarray Data on Matched Frozen Tissues. PLoS ONE. 2016;11: e0153784.doi:10.1371/journal.pone.0153784.

[17] Ragulan C, Eason K, Fontana E, Nyamundanda G, Tarazona N, Patil Y, et al. Analytical Validation of Multiplex Biomarker Assay to Stratify Colorectal Cancer into Molecular Subtypes. Sci Rep. 2019;9:7665. 10.1038/s41598-019-43492-02019.

[18] Guinney J, Dienstmann R, Wang X, de Reyniès A, Schlicker A, Soneson C, et al. The consensus molecular subtypes of colorectal cancer. Nature Med. 2015;21:1350 – 1356.

[19] Xu C, Xia P, Li J, Lewis KB, Ciombor KK, Wang Let al. Discovery an validation of a 10-gene predicitve signature for response to adjuvant chemotherapy in stage II and III colon cancer. Cell Rep Med. 2024;5:e101661. 10.1016/j.xcrm.2024.101661.

[20] Yin H, Harrison TA, Thomas SS, Sather CL, Koehne AL, Malen RC, et al. T cell-inflamed gene expression profile is associated with favorable disease-specific survival in non-hypermutated microsatellite-stable colorectal cancer patients. Cancer Med. 2023;12:6583 – 6593.

